# CountESS: a flexible, graphical pipeline tool for deep mutational scanning analysis

**DOI:** 10.64898/2026.04.27.721230

**Authors:** Nick Moore, Callum J Sargeant, Matthew J Wakefield, Nicholas A Popp, Douglas M Fowler, Alan F Rubin

## Abstract

Deep Mutational Scanning (DMS) experiments generate large volumes of sequencing data that must be processed through multi-step computational pipelines to yield interpretable variant scores. At least twelve dedicated tools have been published for this purpose, yet the diversity of experimental designs, scoring strategies, and software implementations has produced a fragmented landscape in which no single tool accommodates the full range of workflows encountered in practice. Here we present CountESS (Count-based Experiment Scoring and Statistics), an open-source pipeline tool that provides a modular, graphical interface for constructing flexible DMS analysis workflows. CountESS supports a wide range of input formats, barcode translation, HGVS variant calling, and user-defined scoring functions, enabling it to accommodate diverse experimental designs including selection assays, time-series experiments, and bin-based assays such as VAMP-seq. Implemented in Python with DuckDB as a computational backend, the software provides high-performance, memory-efficient processing suitable for large datasets. CountESS is freely available at https://github.com/CountESS-Project/CountESS under the 3-Clause BSD Licence. Supplementary data, including demonstration pipelines and example datasets, are available at https://github.com/CountESS-Project/countess-demo.

## Introduction

Deep Mutational Scanning (DMS) is an experimental technology designed to systematically measure many thousands of protein variant effects in a pooled assay (Fowler and Fields 2014). DMS has been applied broadly for understanding protein function and stability (Olson, Wu, and Sun 2014, Matreyek *et al*. 2018), elucidating protein evolution (Hietpas, Jensen, and Bolon 2011, Firnberg *et al*. 2014, Starr, Picton, and Thornton 2017), providing evidence for clinical variant classification (Findlay *et al*. 2018, Loggerenberg van *et al*. 2023, McEwen *et al*. 2025, Tejura *et al*. 2026), benchmarking predictive models (Frazer *et al*. 2021, Livesey and Marsh 2023), and examining host-pathogen interactions (Thyagarajan and Bloom 2014, Haddox *et al*. 2018, Dadonaite *et al*. 2024).

A typical DMS analysis pipeline involves several discrete computational steps: reading raw sequencing data, counting variant occurrences across experimental conditions, normalising counts, and computing enrichment scores. In its simplest form, a variant effect score is calculated as the log-ratio of a variant’s frequency in a post-selection sample to its frequency in a pre-selection input sample (Fowler *et al*. 2011). However, the diversity of DMS experimental designs substantially complicates this picture (Çubuk *et al*. 2025). Pooled assays may measure cell growth, protein binding, enzymatic activity, or fluorescence, and may be structured as two-population experiments, time-series experiments tracking variant frequencies across multiple time points, or bin-based experiments in which populations are sorted into discrete bins by fluorescence-activated cell sorting (FACS). Sequencing strategies are equally varied, encompassing single-end direct sequencing, overlapping paired-end sequencing, tile-based sequencing, and barcode-based sequencing in which short barcodes must be mapped to full-length variant sequences via a separately generated barcode-variant table.

At least twelve dedicated computational tools have been published for DMS data analysis, including Enrich (Fowler *et al*. 2011), Enrich2 (Rubin *et al*. 2017), DiMSum (Faure *et al*. 2020), mutscan (Soneson *et al*. 2023), Rosace(Rao *et al*. 2024), and gyōza (Durand, Pageau, and Landry 2026), among others. These tools differ substantially in their statistical models, supported experimental designs, input and output formats, and software implementations (Çubuk *et al*. 2025). Most are designed around a fixed analytical workflow suited to a particular experimental paradigm: for example, DiMSum is designed exclusively for two-population experiments, dms_tools2 (Bloom 2015) and dms_variants are optimised for viral barcode-based datasets, and the only dedicated tool that currently exists for bin-based FACS assays (Freudenberg *et al*. 2026), which differs markedly from the way most bin-based experiments are analysed using custom scripts(Matreyek *et al*. 2018). The lack of standardisation in input files and output formats creates a significant barrier for researchers who wish to compare approaches or adapt an existing pipeline to a new experimental design. Furthermore, several tools are no longer actively maintained, creating reproducibility concerns as software dependencies evolve.

Here we describe CountESS (Count-based Experiment Scoring and Statistics), a new open-source tool that addresses these limitations through a modular pipeline architecture. Rather than implementing a fixed analytical workflow, CountESS provides a library of composable processing nodes (plugins) that users assemble into a directed acyclic graph via a graphical user interface (GUI). This design makes CountESS adaptable to the full diversity of DMS experimental designs, including two-population, time-series, and bin-based assays, without requiring users to write bespoke analysis scripts. This flexible approach enables compatibility with emerging and forthcoming experimental designs, and enables rapid prototyping and innovation for bioinformatics methods developers.

## Features and Implementation

### Architecture and Interface

CountESS is implemented in Python and uses DuckDB as its core data processing engine (Raasveldt and Muehleisen, n.d.). DuckDB is an in-process analytical database optimised for columnar, vectorised query execution. It parallelises automatically across available CPU cores and processes data in a streaming fashion, enabling CountESS to handle datasets that exceed available RAM without requiring explicit batching by the user. This provides substantial performance and usability improvements over other data frame based approaches (e.g. pandas (McKinney 2010)), which are memory-bound and single-threaded for most operations. This gives CountESS substantially better performance even on small datasets (Figure 1).

**Figure 1.**
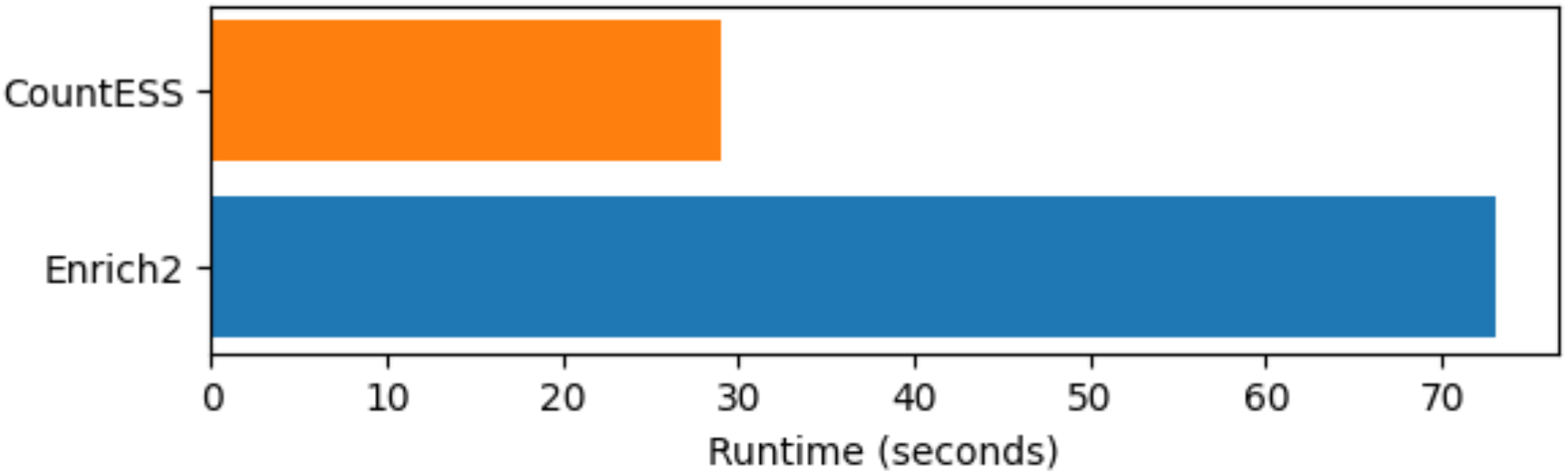
CountESS runtime comparison. CountESS runs substantially faster than Enrich2 on the same dataset from the Enrich2 example repository (https://github.com/FowlerLab/Enrich2-Example). This is due to improved performance from DuckDB as well as parallelisation of sequence file reading steps.

The core architecture treats each analysis step as a plugin that receives one or more relational tables as input, applies a transformation, and passes the result to downstream plugins. Pipelines are defined in human-readable INI configuration files, which can be created and edited either through the GUI or directly in a text editor, supporting both interactive exploration and scripted, reproducible execution. Parallel execution of independent pipeline branches is orchestrated automatically, making efficient use of multi-core hardware.

The GUI is built using Tkinter and provides a canvas on which plugins are arranged and connected (Figure 2). Each plugin exposes a configuration panel in which parameters are set, and a live data preview is updated as parameters are adjusted, enabling rapid iteration during pipeline development. Full pipeline execution is triggered by a single Run command in the GUI, or by command line invocation either locally or in a high-performance computing environment.

**Figure 2.**
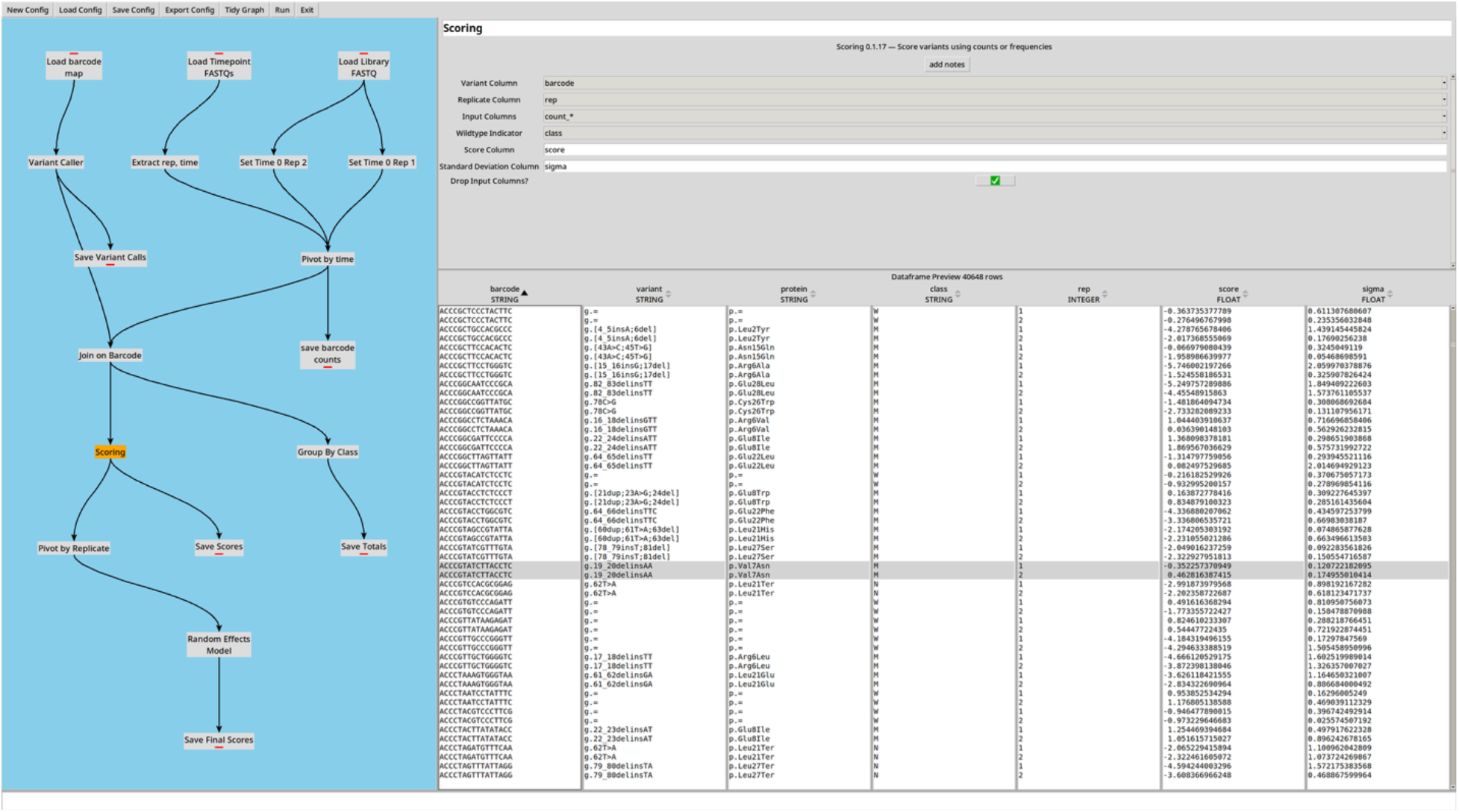
CountESS interface screenshot. The CountESS graphical user interface is split into panes showing the overall workflow graph and associated plugin nodes (left) and the configuration and data frame preview (right). The workflow displayed replicates a barcoded time series analysis from the Enrich2 example repository (https://github.com/FowlerLab/Enrich2-Example).

### Input and Data Handling

CountESS can read sequence data in FASTQ format or load counts and other data from CSV or TSV files, with transparent support for compressed input. Multiple files can be loaded simultaneously and concatenated, and a variety of read filtering options are supported. Raw sequencing reads from input FASTQ files are processed and counted on a per-file basis and can be further aggregated using Group By plugins.

Where experimental metadata (such as time point, replicate, or selection bin) is encoded in the file name rather than file contents, CountESS provides tooling to tokenise file names and extract metadata into named, user-defined columns. For cases where metadata is stored in a separate file (e.g., a run table from the NCBI Sequence Read Archive (Katz *et al*. 2022)), the metadata table can be easily merged with the sequencing data via a Join plugin.

Processed data can be written to CSV or TSV files using the CSV Writer plugin. Because CountESS supports multiple outputs from a single plugin, a single pipeline can simultaneously produce, for example, separate output files for DNA-level and protein-level variant scores without duplicating upstream processing steps. Intermediate files for quality control, visualisation or other purposes are also very straightforward to generate using this system.

### Barcode Translation and Variant Calling

Many DMS libraries are sequenced via short barcode sequences that must be mapped to their corresponding full-length variants (Matreyek *et al*. 2018). CountESS supports this through a Join plugin that merges a barcode-to-sequence mapping file with the counted barcode data. Variant annotation is performed by the Variant Caller plugin, which compares each observed sequence against a user-supplied reference sequence and outputs variant strings in HGVS format (Dunnen den *et al*. 2016) for DNA and/or protein variants. This enables downstream analyses to operate on compact, standardised variant identifiers, and facilitates integration with community databases such as MaveDB (Rubin *et al*. 2025).

### Pivoting and Scoring

To compare variant counts across experimental conditions, CountESS provides a Pivot Tool plugin that reshapes data from long to wide format, expanding a specified column (e.g., time point or selection bin) into separate count columns for each condition. This pivoted representation is the natural input for score calculation.

Variant scoring is performed using either a pre-built plugin that implements an existing statistical method or via the Expression plugin, which allows the user to write arbitrary Python-like expressions that can generate one or more output columns. The latter option provides maximum flexibility and enables rapid prototyping, as the preview data frame updates in real time as the expression is altered, while the former promotes reproducibility and ease for users who are not enaging in statistical methods development. Crucially, because the scoring function can be user-defined rather than hard-coded, CountESS is not limited to any particular statistical model and can accommodate novel scoring approaches as the field develops.

At present, CountESS fully re-implements the core features of Enrich2 (Rubin *et al*. 2017), including log-ratio based scoring, regression-based scoring, and replicate combination using a random effects model (Demidenko 2013). It also includes pre-built plugins for FACS-based scoring as implemented in VAMP-seq (Matreyek *et al*. 2018), with support for custom bin weights.

### Extensibility

CountESS is designed to be extensible. New plugins can be implemented by subclassing a base plugin class and registering the plugin with the CountESS framework, allowing research groups to develop and share custom plugins for specialised processing steps while benefiting from the existing pipeline infrastructure.

While most CountESS plugins are written in Python, the existing classes can be used as wrappers to provide code from other languages, or access to third party tools. For example, the CountESS minimap2 plugin (https://github.com/CountESS-Project/countess-minimap2) allows the user to process sequences using the minimap2 aligner (Li 2018), generating an HGVS variant as well as the underlying CIGAR string and other alignment information.

## Discussion

CountESS addresses a practical need in the DMS community for a flexible, user-friendly analysis tool that does not impose a fixed analytical workflow. The existing landscape of DMS scoring tools is characterised by fragmentation: tools differ in their supported experimental designs, input formats, statistical models, and software implementations, and the lack of standardisation creates a significant cost for researchers who wish to explore alternative analysis strategies or apply existing tools to new experimental designs (Çubuk *et al*. 2025). CountESS takes a complementary approach to this problem. Rather than implementing a new statistical scoring method, it provides a shared computational framework within which diverse scoring strategies, from simple log-ratio enrichment to weighted bin sums to user-defined models, can be constructed, inspected, and executed using a consistent interface.

The node-based graphical interface lowers the barrier to entry for researchers who are not specialist bioinformaticians, while the Expression plugins and INI-based pipeline configuration retain the flexibility required by expert users. The use of DuckDB as a backend ensures that performance scales with dataset size and available hardware, addressing a practical limitation of earlier pandas-based tools for large-scale experiments.

Future work on the project will include built-in implementations of more sophisticated statistical models offered by tools such as Rosace (Rao *et al*. 2024) or DiMSum (Faure *et al*. 2020), as well as a more comprehensive library of example workflows.

## Availability and Implementation

CountESS is implemented in Python and is freely available to non-commercial users under the 3-Clause BSD Licence. Source code, installation instructions, and documentation are available at: https://github.com/CountESS-Project/CountESS

Demonstration pipelines and example datasets are available at https://github.com/CountESS-Project/countess-demo

The software has been tested on Linux, macOS, and Windows.

## Acknowledgements

The authors wish to thank all CountESS and Enrich2 users. Their feedback and experiences helped inform this work. This study was supported by NHGRI IGVF grant UM1HG011969.

